# Clinical Characteristics and Risk of Second Primary Lung Cancer After Cervix Cancer: A Population-Based Study

**DOI:** 10.1101/2020.04.02.021626

**Authors:** Chengyuan Qian, Hong Liu, Yan Feng, Shenglan Meng, Dong Wang, Mingfang Xu

## Abstract

**Background:** Lung cancer as a second primary malignancy is increasingly common, but the clinical characteristics of second primary non-small cell lung cancer after cervix cancer (CC-NSCLC) in comparison with first primary non-small cell lung cancer (NSCLC1) is unknown.

**Methods:** The Surveillance, Epidemiology and EndResults (SEER) cancer registry between 1998 and 2010 was used to conduct a large population-based cohort analysis. Demographic and clinical characteristics as well as prognostic data were systematically analyzed. We further compared overall survival (OS) in the two cohorts. Risk factors of secondary primary lung cancer in cervical cancer patients were also analyzed.

**Results:** 557 (3.52%) had developed second primary lung cancer after cervix cancer and 451 were eligible for inclusion in the final analyses. In comparison to NSCLC1, patients with CC-NSCLC had a higher rate of squamous cell carcinoma (SCC) (36.59% vs. 19.07%, p<0.01). Median OS was longer for CC-NSCLC than for NSCLC1 before propensity score matching (PSM) (16 vs. 13months) but there was no significant difference after PSM. High-risk factors in cervical cancer to developing CC-NSCLC include: 50-79years old, black race (OR 1.417; 95%CI 1.095-1.834; p<0.05)and history of radiotherapy (OR 1.392; 95%CI 1.053-1.841; p<0.05).

**Conclusion:** 50-79years old, black race and history of radiotherapy were independent risk factors of second primary lung cancer in cervical cancer patient. CC-NSCLC patients had distinctive clinical characteristics and a better prognosis compared with NSCLC1 patients.

## Introduction

Cervical cancer is the fourth most common cancer worldwide, with an estimated 310 000 annual deaths globally ^1–3^. However, improvements in early detection and cancer treatment have led to long survival among cervical cancer patients. Subsequently, the possibility for patients to develop a subsequent primary cancer becomes a more important consideration ^4^, with a 17% higher rate of cancer than general population in female. What’s more, cervix cancer survivors had more than double the rate of lung cancer ^5^. As a matter of fact, from 1975 to 2001, 756,467 people in the United States have developed a second solid cancer, representing almost 8 percent of the current cancer survivor population. Subsequent malignancies in cancer survivors now constitute 18% of all cancer diagnosis in the US SEER cancer registries ^4^.

In particular, lung cancer as a second primary malignancy is increasingly common. Indeed, lung cancer is the leading cause of cancer incidence and mortality around the world, with 2.1 million new lung cancer cases and 1.8 million deaths predicted in 2018, representing 18.4% cancer deaths ^1^. Among women, lung cancer constitutes one of the 3 most commonly diagnosed cancers besides breast and colorectal cancers ^2^. Notably, lung cancer incidence rates are now higher among young women than among young men in non-Hispanic whites and Hispanics Americans ^6^. As regards both cervical cancer and lung cancer, approximately 10% of cervical cancer survivors have developed a second malignancy, in which lung cancer accounts for one of the largest numbers ^7,8^. However, risk factors of secondary primary lung cancer in cervical cancer patients are not known. Similarly, differences between CC-NSCLC and NSCLC on clinical characteristics and survival have not been studied. Consequently, there is a crucial need to characterize the lung cancer disease in this specific subgroup as regards both their high incidence rates. Thereby, our study aims to focus on clinical differences between CC-NSCLC and NSCLC1 as well as risk factors for secondary primary lung cancer in patients with cervical cancer.

## Methods

### Ethical Statement

The study was approved by the Research Ethics Committee of Daping Hospital. Data obtained from SEER database did not require informed patient consent because cancer is a reportable disease in the United States.

### Population

We identified cervical cancer cases and NSCLC cases from the SEER program of the National Cancer Institute (http://seer.cancer.gov/). The cohort was composed of adult patients who were pathologically confirmed with cervical cancer or NSCLC from the SEER database from 1998 to 2010. This SEER program released a 18 population-based cancer registries of incidence rate and survival rate in United States, covering about 28% of the general population. Exclusion criteria were: confirmed by autopsy, unknown age of diagnosis, unknown marriage status, undetermined grade of disease, unknown stage of disease, unknown pathological type. A total of 173272 NSCLC1 patients, 15809 cervical cancer patients and 451 CC-NSCLC patients were eligible for inclusion in the final analyses. Domestic status was recorded as follows: never married as “unmarried”, married as married or unmarried but having domestic partner; separated, divorced and widowed status were classified as “other”. Except for squamous cell neoplasm and adenocarcinoma, other histology types were recorded as “other”, including NSCLC not otherwise specified (NSCLC-NOS).

### Statistical analysis

Categorical measurements were described as count and percentage, while continuous measurements were presented as mean (median) and range. The chi-square was used to compare the categorical measurements while the t test was used for continuous ones. Survival data were measured from the lung cancer date of diagnosis to the date of all-cause death or the last follow-up. Cumulative survival curves were generated by the Kaplan-Meier method. Differences in survival were compared using the log-rank (Mantel-Cox) tests. PSM was used to balance the difference from baseline characteristics between CC-NSCLC and NSCLC1 groups. According to 1 to 3 matches, 449 CC-NSCLC patients were matched successfully. After PSM, there were 449 cases in the CC-NSCLC group and 1347 cases in the NSCLC1 group, and there were no significant differences in histology, age at lung cancer diagnosis, race, year of lung cancer diagnosis, stage of lung cancer, marital status, radiotherapy records, chemotherapy records, surgery records and grade between the two groups. Logistic multiple regression analysis was performed to identify independent risk factors and odds ratios (OR) of second primary lung cancer in cervical cancer patient. All p values were two-sided, with p < 0.05 considered statistically significant. The incidence of second primary lung cancer was compared to the literature. All the analyses were done using SPSS statistical software, version 23 (IBM Corp, Armonk, NY).

## Results

### Patients Characteristics

A total of 15809 cervical cancer patients and 173272 NSCLC1 patients between 1998 and 2010 were involved. 557 patients (3.52%) were diagnosed with second primary non-small cell lung cancer after cervix cancer (CC-NSCLC) and 451 patients with complete information were eligible for inclusion in the final analyses. The demographic and clinic-pathologic features of NSCLC1 and CC-NSCLC patients are listed in Table 1. There are significant differences in histology, age at diagnosis, race, year at diagnosis, marital and cause of death between CC-NSCLC and NSCLC1. No significant was detected in stage, radiotherapy records, chemotherapy records, surgery records and grade. The mean time to NSCLC diagnosis was 57 months after cervical cancer, with a range of 12–192 months. The mean age of cervical cancer diagnosis was 58.2 years whereas the mean age at CC-NSCLC diagnosis was 62.9 years. The majority of CC-NSCLC were adenocarcinomas (38.36%) while 36.59% were SCC and 25.1% were other. Of the 173272 NSCLC1 patients in the database, a vast majority was adenocarcinomas (49.05%), and 19.07% of patients were SCC while 31.89% of patients were other. The proportion of SCC in CC-NSCLC patients was apparently higher than that in NSCLC1 patients (36.59% vs. 19.07%). The difference in pathologic type distribution between these two cohorts is significant (p<0.01).

**Table 1.**
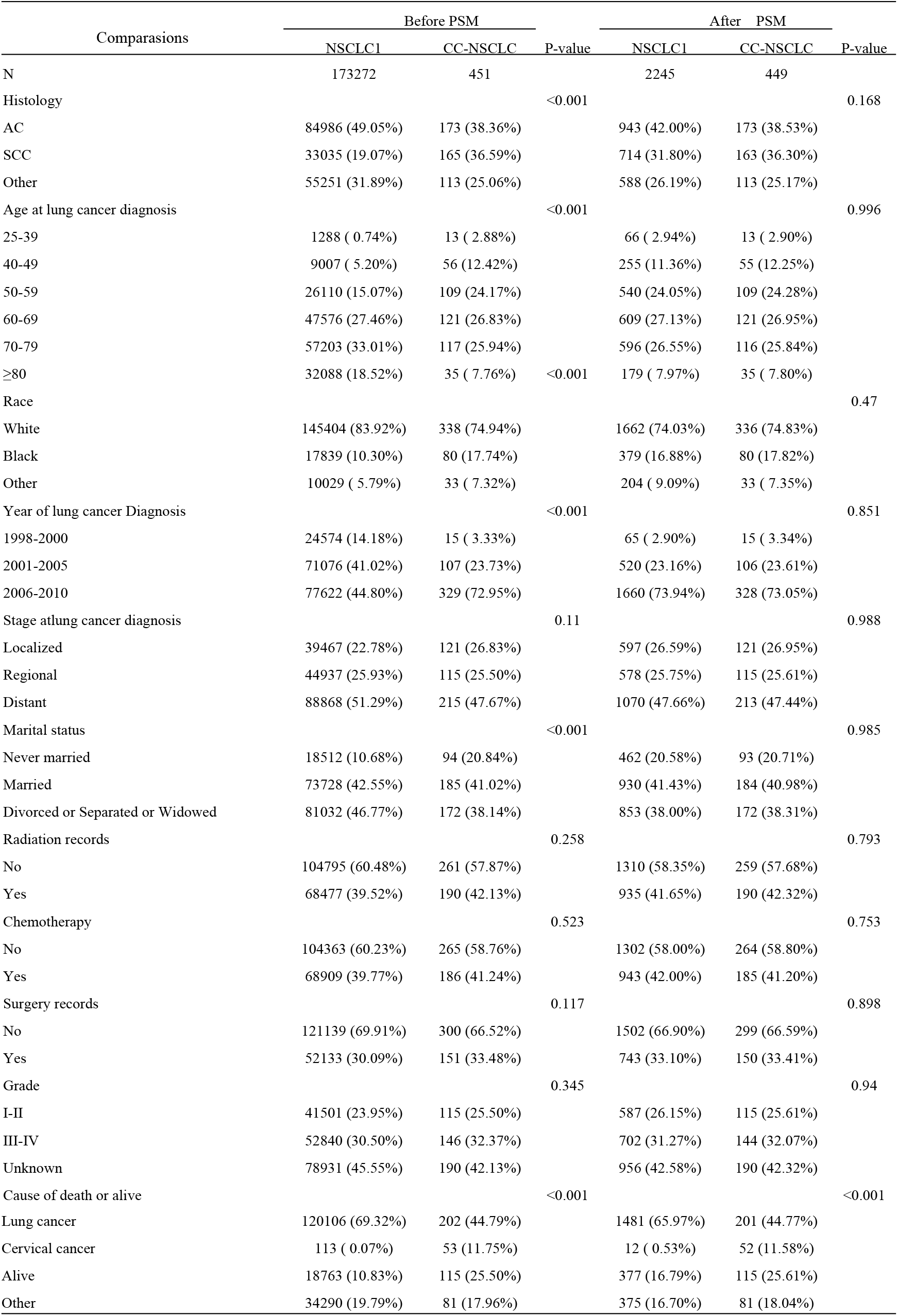
Demographic and clinic-pathological characteristics of patients with CC-NSCLC and NSCLC before and after PSM

### Clinical features in CC-NSCLC patients

The impact of demographic characteristics and clinical features of cervical cancer on pathological types and clinical stages of lung cancer in CC-NSCLC patients are listed in Table 2. Latency, stage, histology, radiotherapy records, chemotherapy records and grade of cervical cancer were associated with pathological types of lung cancer, rather than race, age at cervical cancer diagnosis, year of cervical cancer diagnosis, marital status, surgery records. There was no significant correlation between clinical factors of cervical cancer and stages of lung cancer in CC-NSCLC patients.

**Table 2.** Impact of demographic and clinical features of cervical cancer on pathological and stage of lung cancer

| Cervical cancers | Lung cancers |  |  |  | P value | Lung cancers |  |  | P-value |
| --- | --- | --- | --- | --- | --- | --- | --- | --- | --- |
|  | Total | AC(%) | SCC(%) | Other(%) |  | Localized(%) | Regional(%) | Distant(%) |  |
| Total | 451 | 173 (38.4) | 165 (36.6) | 113 (25.1) |  | 121 (26.8) | 115 (25.5) | 215 (47.7) |  |
| Latency |  |  |  |  | <0.05 |  |  |  | >0.05 |
| ≤1year | 100 | 42 (42.0) | 31 (31.0) | 27 (27.0) |  | 37 (37.0) | 23 (23.0) | 40 (40.0) |  |
| >1year, ≤5years | 187 | 57 (30.5) | 69 (36.9) | 61 (32.6) |  | 43 (23.0) | 48 (25.7) | 96 (51.3) |  |
| >5years, ≤10years | 117 | 51 (43.6) | 47 (40.2) | 19 (16.2) |  | 27 (23.1) | 34 (29.0) | 56 (47.9) |  |
| >10years | 47 | 23 (48.9) | 18 (38.3) | 6 (12.8) |  | 14 (29.8) | 10 (21.3) | 23 (48.9) |  |
| Race |  |  |  |  | >0.05 |  |  |  | >0.05 |
| White | 338 | 127 (37.6) | 122 (36.1) | 89 (26.3) |  | 97 (28.7) | 84 (24.9) | 157 (46.4) |  |
| Black | 80 | 29 (36.3) | 36 (45.0) | 15 (18.8) |  | 16 (20.0) | 23 (28.7) | 41 (51.3) |  |
| Other | 33 | 17 (51.5) | 7 (21.2) | 9 (27.3) |  | 8 (24.2) | 8 (24.2) | 17 (51.6) |  |
| Age at cervical cancer diagnosis |  |  |  |  | >0.05 |  |  |  | >0.05 |
| 25-39 | 35 | 13 (37.1) | 15 (42.9) | 7 (20.2) |  | 8 (22.9) | 10 (28.6) | 17 (48.6) |  |
| 40-49 | 77 | 20 (26.0) | 33 (42.9) | 24 (31.1) |  | 16 (20.8) | 19 (24.7) | 42 (54.5) |  |
| 50-59 | 133 | 61 (45.9) | 39 (29.3) | 33 (24.8) |  | 32 (24.1) | 35 (26.3) | 66 (49.6) |  |
| 60-69 | 124 | 53 (42.7) | 43 (34.7) | 28 (22.6) |  | 39 (31.5) | 33 (26.6) | 52 (41.9) |  |
| 70-79 | 68 | 21 (30.9) | 28 (41.2) | 19 (27.9) |  | 20 (29.4) | 17 (25.0) | 31 (45.6) |  |
| ≥80 | 14 | 5 (35.7) | 7 (50.5) | 2 (14.3) |  | 6 (42.9) | 1 (7.1) | 7 (50.0) |  |
| Year of cervical cancer Diagnosis |  |  |  |  | >0.05 |  |  |  | >0.05 |
| 1998-2000 | 112 | 36 (32.1) | 43 (38.4) | 33 (29.5) |  | 28 (25.0) | 30 (26.8) | 54 (48.2) |  |
| 2001-2005 | 216 | 84 (38.9) | 81 (37.5) | 51 (23.6) |  | 62 (28.7) | 51 (23.6) | 103 (47.7) |  |
| 2006-2010 | 123 | 53 (43.1) | 41 (33.3) | 29 (23.6) |  | 31 (25.2) | 34 (27.6) | 58 (47.2) |  |
| Stage at CC cancer diagnosis |  |  |  |  | <0.001 |  |  |  | >0.05 |
| localized | 213 | 103 (48.4) | 61 (28.6) | 49 (23.0) |  | 60 (28.2) | 57 (26.8) | 96 (45.0) |  |
| regional | 238 | 70 (29.4) | 104 (43.7) | 64 (26.9) |  | 61 (25.6) | 58 (24.4) | 119 (50.0) |  |
| Histology |  |  |  |  | p<0.001 |  |  |  | >0.05 |
| AC | 89 | 49 (55.0) | 15 (16.9) | 25 (28.1) |  | 20 (22.5) | 24 (27.0) | 45 (50.5) |  |
| SCC | 335 | 111 (33.1) | 146 (43.6) | 78 (23.3) |  | 94 (28.1) | 85 (25.4) | 156 (46.6) |  |
| other | 27 | 13 (48.2) | 4 (14.8) | 10 (37.0) |  | 7 (25.9) | 6 (22.2) | 14 (51.9) |  |
| Marital status |  |  |  |  | >0.05 |  |  |  | >0.05 |
| Never married | 94 | 33 (35.1) | 38 (40.4) | 23 (24.5) |  | 19 (20.2) | 24 (25.5) | 51 (54.3) |  |
| Married | 185 | 82 (44.3) | 57 (30.8) | 46 (24.9) |  | 50 (27.0) | 48 (25.9) | 87 (47.1) |  |
| Divorced or Separated or Widowed | 172 | 58 (33.7) | 70 (40.7) | 44 (25.6) |  | 52 (30.2) | 43 (25.0) | 77 (44.8) |  |
| Radiation records |  |  |  |  | <0.001 |  |  |  | >0.05 |
| No | 151 | 78 (51.7) | 37 (24.5) | 36 (23.8) |  | 82 (27.3) | 70 (23.3) | 148 (49.4) |  |
| Yes | 300 | 95 (31.7) | 128 (42.7) | 77 (25.6) |  | 39 (25.8) | 45 (29.8) | 67 (44.4) |  |
| Chemotherapy records |  |  |  |  | <0.001 |  |  |  | >0.05 |
| No | 247 | 117 (47.4) | 74 (30.0) | 56 (22.6) |  | 55 (27.0) | 46 (22.5) | 103 (50.5) |  |
| Yes | 204 | 56 (27.5) | 91 (44.6) | 57 (27.9) |  | 66 (26.7) | 69 (27.9) | 112 (45.3) |  |
| Surgery records |  |  |  |  |  |  |  |  | >0.05 |
| No | 194 | 56 (28.9) | 87 (44.8) | 51 (26.3) |  | 69 (26.8) | 63 (24.5) | 125 (48.7) |  |
| Yes | 257 | 117 (45.5) | 78 (30.4) | 62 (24.1) |  | 52 (26.8) | 52 (26.8) | 90 (46.4) |  |
| Grade |  |  |  |  | <0.05 |  |  |  | >0.05 |
| I-II | 180 | 77 (42.8) | 69 (38.3) | 34 (18.9) |  | 55 (30.6) | 48 (26.7) | 77 (42.8) |  |
| III-IV | 162 | 51 (31.5) | 64 (39.5) | 47 (29.0) |  | 33 (20.4) | 37 (22.8) | 92 (56.8) |  |
| Unknown | 109 | 45 (41.3) | 32 (29.4) | 32 (29.4) |  | 33 (30.3) | 30 (27.5) | 46 (42.2) |  |

The impact of demographic characteristics and clinical features of cervical cancer on the cause of death in CC-NSCLC patients are listed in Table 3. Latency, age at diagnosis, stage, histology, marital status, radiotherapy records, chemotherapy records, surgery records and grade were associated with the causes of death, rather than race and year of cervical cancer diagnosis. Patients with a latency≤1year were more likely to die of cervical cancer, and those with a latency>5years were more likely to survive. Married patients with young age, regional stage, treated by radiation or chemotherapy, died more often from cervical cancer. Patients with cervical adenocarcinoma, well or moderately differentiated in terms of histological grade and treated by surgery were more likely to survive. Lung cancer was the most common cause of death (44.8%).

**Table 3.** Impact of demographic and clinical features of cervical cancer on the death in CC-NSCLC patients

| Cervical cancers | Cause of death or alive |  |  |  |  | P-value |
| --- | --- | --- | --- | --- | --- | --- |
|  | Total | Lung and Bronchus(%) | Cervix uteri(%) | Alive(%) | Other(%) |  |
| Total | 451 | 202 (44.8) | 53 (11.8) | 115 (25.5) | 81 (18.0) |  |
| Latency |  |  |  |  |  | <0.001 |
| ≤1year | 100 | 42 (42.0) | 18 (18.0) | 12 (12.0) | 28 (28.0) |  |
| >1year, ≤5years | 187 | 86 (46.0) | 24 (12.8) | 42 (22.5) | 35 (18.7) |  |
| >5years, ≤10years | 117 | 54 (46.2) | 9 (7.7) | 40 (34.2) | 14 (12.0) |  |
| >10years | 47 | 20 (42.6) | 2 (4.3) | 21 (44.7) | 4 (8.5) |  |
| Race |  |  |  |  |  | >0.05 |
| White | 338 | 153 (45.3) | 43 (12.7) | 85 (25.1) | 57 (16.9) |  |
| Black | 80 | 35 (43.8) | 8 (10.0) | 18 (22.5) | 19 (23.8) |  |
| Other | 33 | 14 (42.4) | 2 (6.1) | 12 (36.4) | 5 (15.2) |  |
| Age at cervical cancer diagnosis |  |  |  |  |  | <0.001 |
| 25-39 | 35 | 7 (20.0) | 10 (28.6) | 15 (42.9) | 3 (8.5) |  |
| 40-49 | 77 | 35 (45.5) | 11 (14.3) | 19 (24.7) | 12 (15.5) |  |
| 50-59 | 133 | 61 (45.9) | 16 (12.0) | 38 (28.6) | 18 (13.5) |  |
| 60-69 | 124 | 55 (44.4) | 11 (8.9) | 33 (26.6) | 25 (20.1) |  |
| 70-79 | 68 | 37 (54.4) | 3 (4.4) | 8 (11.8) | 20 (29.4) |  |
| ≥80 | 14 | 7 (50.0) | 2 (14.3) | 2 (14.3) | 3 (21.4) |  |
| Year of cervical cancer diagnosis |  |  |  |  |  | >0.05 |
| 1998-2000 | 112 | 51 (45.5) | 14 (12.5) | 26 (23.2) | 21 (18.8) |  |
| 2001-2005 | 216 | 104 (48.1) | 27 (12.5) | 55 (25.5) | 30 (13.9) |  |
| 2006-2010 | 123 | 47 (38.2) | 12 (9.8) | 34 (27.6) | 30 (24.4) |  |
| Stage at breast cancer diagnosis |  |  |  |  |  | <0.05 |
| Localized | 213 | 96 (45.1) | 12 (5.6) | 67 (31.5) | 38 (17.8) |  |
| Regional | 238 | 106 (44.5) | 41 (17.2) | 48 (20.2) | 43 (18.1) |  |
| Histology |  |  |  |  |  | p<0.05 |
| AC | 89 | 32 (36.0) | 7 (7.9) | 33 (37.1) | 17 (19.0) |  |
| SCC | 335 | 155 (46.3) | 41 (12.2) | 80 (23.9) | 59 (17.6) |  |
| Other | 27 | 15 (55.6) | 5 (18.5) | 2 (7.4) | 5 (18.5) |  |
| Marital status |  |  |  |  |  | <0.05 |
| Never married | 94 | 44 (46.8) | 7 (7.4) | 26 (27.7) | 17 (18.1) |  |
| Married | 185 | 75 (40.5) | 29 (15.7) | 56 (30.3) | 25 (13.5) |  |
| Divorced or Separated or Widowed | 172 | 83 (48.3) | 17 (9.9) | 33 (19.2) | 39 (22.7) |  |
| Radiation records |  |  |  |  |  | <0.05 |
| Yes | 300 | 134 (44.7) | 46 (15.3) | 64 (21.3) | 56 (18.7) |  |
| No | 151 | 68 (45.0) | 7 (4.6) | 51 (33.8) | 25 (16.6) |  |
| Chemotherapy records |  |  |  |  |  | <0.05 |
| Yes | 204 | 85 (41.7) | 37 (18.1) | 47 (23.0) | 35 (17.2) |  |
| No | 247 | 117 (47.4) | 16 (6.5) | 68 (27.5) | 46 (18.6) |  |
| Surgery records |  |  |  |  |  | <0.05 |
| Yes | 257 | 110 (42.8) | 24 (9.3) | 83 (32.3) | 40 (15.6) |  |
| No | 194 | 92 (47.4) | 29 (14.9) | 32 (16.5) | 41 (21.1) |  |
| Grade |  |  |  |  |  | <0.05 |
| I-II | 180 | 73 (40.6) | 18 (10.0) | 60 (33.3) | 29 (16.1) |  |
| III-IV | 162 | 82 (50.6) | 24 (14.8) | 32 (19.8) | 24 (14.8) |  |
| Unknown | 109 | 47 (43.1) | 11 (10.1) | 23 (21.1) | 28 (25.7) |  |

### Risk factors of secondary primary lung cancer in cervical cancer patients

High-risk factors of developing secondary primary lung cancer in cervical cancer patients include: age between 50 and 79years old, black race and history of radiotherapy. All significant independent factors from logistic multiple regression analysis to develop second primary lung cancer are shown in Table 4.

**Table 4.** Risk factors of secondary primary lung cancer in cervical cancer patients

| Comparasions | Cervical cancer | CC-NSCLC | OR (95% CI) | P-value |
| --- | --- | --- | --- | --- |
| Histology |  |  |  | >0.05 |
| AC | 3705 | 89 (19.73%) |  |  |
| SCC | 10506 | 335 (74.28%) | 1.137(0.890-1.454) |  |
| Other | 1147 | 27 ( 5.99%) | 0.832(0.533-1.297) |  |
| Age |  |  |  | <0.05 |
| 25-39 | 5101 (33.21%) | 35 ( 7.76%) |  |  |
| 40-49 | 4413 (28.73%) | 77 (17.07%) | 2.410(1.609-3.609) |  |
| 50-59 | 2680 (17.45%) | 133 (29.49%) | 6.458(4.398-9.483) |  |
| 60-69 | 1588 (10.34%) | 124 (27.49%) | 9.956(6.712-14.767) |  |
| 70-79 | 993 ( 6.47%) | 68 (15.08%) | 8.464(5.470-13.099) |  |
| ≥80 | 583 ( 3.80%) | 14 ( 3.10%) | 2.761(1.427-5.342) |  |
| Race |  |  |  | <0.05 |
| White | 11492 (74.83%) | 338 (74.94%) |  |  |
| Black | 1716 (11.17%) | 80 (17.74%) | 1.417(1.095-1.834) |  |
| Other | 2150 (14.00%) | 33 ( 7.32%) | 0.432(0.300-0.622) |  |
| Year |  |  |  | >0.05 |
| 1998-2000 | 4008 (26.10%) | 112 (24.83%) |  |  |
| 2001-2005 | 6224 (40.53%) | 216 (47.89%) | 1.177(0.929-1.492) |  |
| 2006-2010 | 5126 (33.38%) | 123 (27.27%) | 0.819(0.627-1.069) |  |
| Stage |  |  |  | >0.05 |
| Localized | 9123 (59.40%) | 213 (47.23%) |  |  |
| Regional | 6235 (40.60%) | 238 (52.77%) | 0.877(0.684-1.124) |  |
| Marital status |  |  |  | >0.05 |
| Unmarried | 4480 (29.17%) | 94 (20.84%) |  |  |
| Married | 7371 (47.99%) | 185 (41.02%) | 1.165(0.898-1.510) |  |
| Divorced or Separated or Widowed | 3507 (22.84%) | 172 (38.14%) | 1.391(1.062-1.822) |  |
| Surgery records |  |  |  | >0.05 |
| No | 4687 (30.52%) | 194 (43.02%) |  |  |
| Yes | 10671 (69.48%) | 257 (56.98%) | 0.998(0.768-1.267) |  |
| Radiation records |  |  |  | <0.05 |
| No | 7820 (50.92%) | 151 (33.48%) |  |  |
| Yes | 7538 (49.08%) | 300 (66.52%) | 1.392(1.053-1.841) |  |
| Chemotherapy records |  |  |  | >0.05 |
| No | 9904 (64.49%) | 247 (54.77%) |  |  |
| Yes | 5454 (35.51%) | 204 (45.23%) | 0.989(0.764-1.280) |  |
| Grade |  |  |  | >0.05 |
| I-II | 6100 (39.72%) | 180 (39.91%) |  |  |
| III-IV | 4485 (29.20%) | 162 (35.92%) | 1.057(0.847-1.319) |  |
| Unknown | 4773 (31.08%) | 109 (24.17%) | 0.814(0.634-1.045) |  |

### Survival Analysis

The median OS was 16months (range, 1-191 months) vs. 13months (range, 1-227 months) in CC-NSCLC patients and NSCLC1 patients before PSM, whereas 16months vs. 17months after PSM. The difference was significant before PSM (p < 0.05) but no significant after PSM (p>0.05). Figure1 shows the survival curves of CC-NSCLC and NSCLC1 before and after PSM. OS was longer for CC-NSCLC vs. NSCLC1 before PSM but no significant difference after PSM.

**Figure 1.**
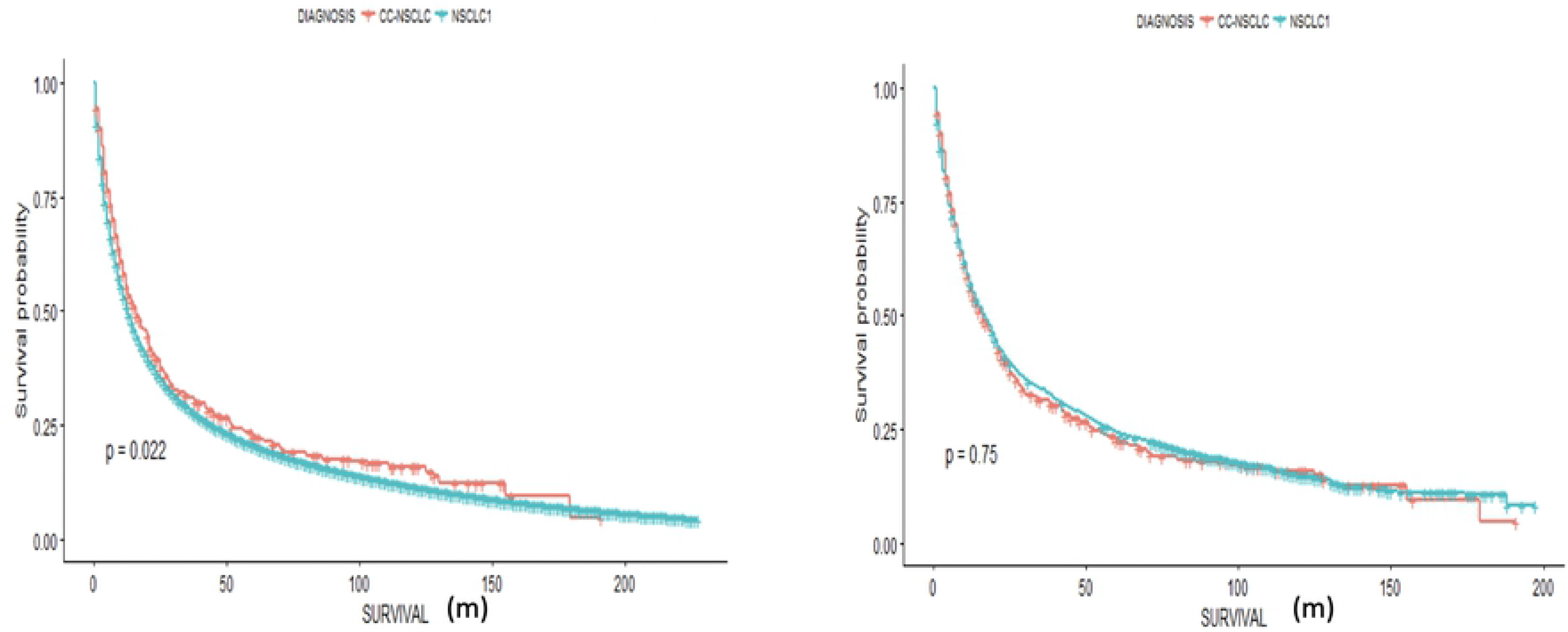
mOS for CC-NSCLC and NSCLC1 before and after PSM. Left, Kaplan Meier survival curves indicated that patients with CC-NSCLC had significantly longer survival than NSCLC1 patients before PSM (16 vs. 13months; p<0.05). Right, Kaplan-Meier survival curves indicated that patients with CC-NSCLC had no significant extension in survival relative to NSCLC1 patients after PSM (16 vs. 15months; p>0.05).

Stage was strongly associated with OS. Median OS of localized, regional and distant CC-NSCLC was 52.0, 25.0, and 8.0m (p < 0.0001). Pathological type was another important prognostic factor for OS. Unsurprisingly, patients with adenocarcinoma had much longer OS than those with SCC or other (22.0, 16.0,and 11.0m respectively (p < 0.01)). Young patients had superior OS in comparison with patients older than 80. The latter had a median OS of 7 months. Differentiation was also an important prognostic factor. mOS of CC-NSCLC patients with grade I+II, grade III+IV and unknown differentiation were respectively 45, 13 and 10m (p < 0.01).Patients who had surgery had a better prognosis (70 vs. 10m, p < 0.01, and those who had radiation had a worse prognosis (13 vs. 121m, p < 0.01) (Figure2).

**Figure 2.**
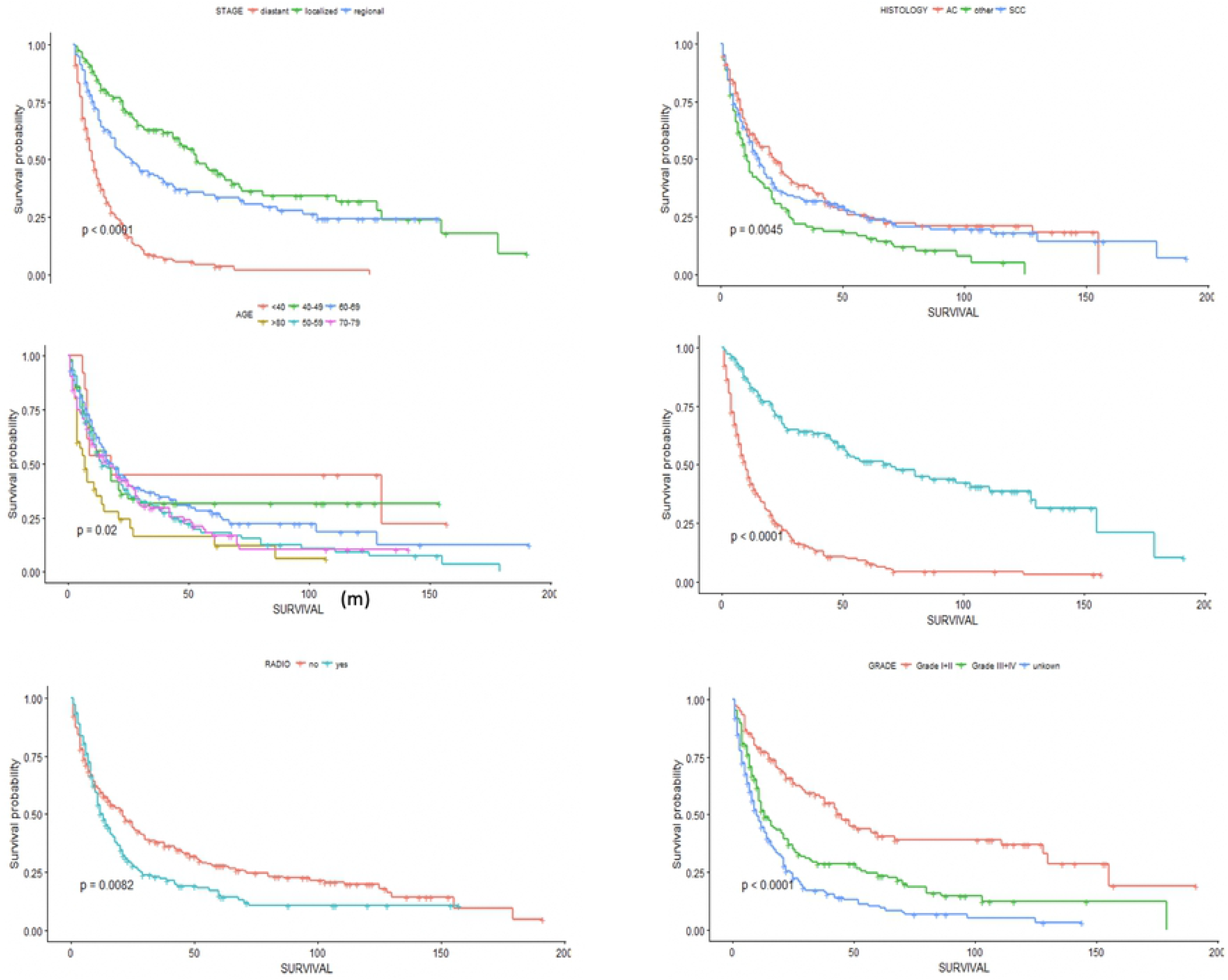
Influence of stage, pathological type, age, surgical records, radiotherapy records and differentiation grade on OS in CC-NSCLC. Kaplan-Meier survival curves indicated that the mOS of CC-NSCLC patients with localized, regional, and distant staging were 52.0, 25.0, and 8.0 months, respectively (p < 0.0001); Kaplan-Meier survival curves indicated that the mOS of CC-NSCLC patients with adenocarcinoma, SCC and other were 22.0, 16.0, and 11.0 months, respectively (p < 0.01); Kaplan-Meier survival curves indicated that the mOS of CC-NSCLC in young patients had superior OS in comparison with patients older than 80 (p < 0.05); Kaplan-Meier survival curves indicated that the mOS of CC-NSCLC patients underwent surgical resection had significantly longer survival than those did not undergo resection (70 vs. 10 months; P<0.0001);Kaplan-Meier survival curves indicated that the mOS of CC-NSCLC patients underwent radiotherapy had significantly shorter survival than those did not undergo radiotherapy(13 vs. 21 months; P<0.01); Kaplan-Meier survival curves indicated that the mOS of CC-NSCLC patients with grade I+II, grade III+IV and unknown differentiation were respectively 45, 13 and 10 months (p < 0.0001).

### Discussion

Over the past three decades, advances in early detection and treatment of cervix cancer have resulted in significant survival improvement among cervical cancer patients. Such survival after a cervical cancer diagnosis are higher than they have ever been, due to improvements in cancer therapy and current emerging issues concern long-term events in survivors, with notably the occurrence of second cancer^9^. Patients will develop subsequent primary cancers as a result of shared lifestyle and genetic factors, as well as the first cancer treatment. In particular, lung cancer accounts for one of the largest numbers in cervical cancer survivors who had developed a second malignancy^7,8^. However, risk factors for secondary primary lung cancer in patients with cervical cancer are poorly known. Similarly, it is unknown if the clinical features of cervical cancer have an impact on the pathological types and stages of secondary lung cancer. Furthermore survival data are lacking as regards differences in terms of prognosis between NSCLC1 and CC-NSCLC. To the best of our knowledge, our study is the first to focus on CC-NSCLC, with the aim to objective differences between CC-NSCLC and NSCLC1 besides the goal to identify risk factors for secondary primary lung cancer in patients with cervical cancer.

Subsequently, our article provides answers to these various issues. First and foremost, CC-NSCLC patients are younger with earlier stages. The proportion of SCC in CC-NSCLC patients was significantly higher than in NSCLC1 patients (36.59% vs. 19.07%). If CC-NSCLC patients seemed to have a better prognosis, no significant difference was found after PSM. Cervical cancer patients who developed secondary primary lung cancer had several characteristics including 50-79years old, black race, and history of radiotherapy.

Concerning epidemiologic data, the incidence of lung cancer differs according to geographical region and over time. In particular, both incidence and mortality from lung cancer continue to increase sharply in China ^10,11^. Our results suggest that the incidence of lung cancer among cervical cancer survivors in our cohort (3.52%) was significantly higher in comparison to rates reported in the literature. According to the region-specific incidence Age-Standardized Rates by Sex for Cancers of the Lung in 2018, female in Northern American have the highest incidence, which is 30.7 per 100,000^1^.

In our cohort, CC-NSCLC patients were younger than NSCLC1 patients, and displayed earlier stages, which may be due to more frequent medical examinations. Squamous cell lung carcinoma was the most common histologic subtype before the 1990s. Currently, adenocarcinoma has become the most common histologic subtype of lung cancer in men and women ^12,13^. Although adenocarcinoma (38.36%) remains the most common histologic subtype in CC-NSCLC patients, the proportion of SCC is significantly higher among NSCLC1 patients (36.59% vs. 19.07%). SCC accounts for 19.07% of NSCLC1 patients in our study similarly to data from literature ^14^. The high proportion of SCC in cervical cancer patients may be due to the history of chemo-radiation therapy as well as confounding factors such as unclear primary and cervical cancer metastasis. Besides, it has to be noted that since 2006, the incidence of second primary lung cancer among cervical cancer patients has drastically increased in our study, which cannot be entirely explained by the increase of lung cancer rates. Indeed, more reasons are required to explain the increased incidence of second primary lung cancer in cervical cancer patients. Interestingly, there was no difference in the number of patients undergoing surgery, chemotherapy and radiotherapy between the two groups. The cancer-related death rate of CC-NSCLC patients was significantly lower than in NSCLC1 patients (56.54% vs. 69.39%), and the survival rate was significantly higher in CC-NSCLC patients than in NSCLC1 patients (25.50% vs. 10.83%), with longer OS for CC-NSCLC patients. However, there was no significant difference in prognosis after PSM between CC-NSCLC and NSCLC1. We infer that the better prognosis of CC-NSCLC before PSM may be due to earlier stages and the young age of the patients.Patients with surgical records, without radiotherapy records had a significant better prognosis. This phenomenon can be partly explained by the fact that more patients in the early stage receive surgical treatment and more patients in the late stage receive radiotherapy. Therefore, cervical cancer patients with high risk factors including 50-79years old, black race, and history of radiotherapy should be reexamined with chest CT scan more frequently for early diagnosis of second primary lung cancer.

As regards the impact of cervical cancer patients clinical features on pathological types of lung cancer, our study highlight that the incidence of SCC is higher in blacks than whites in CC-NSCLC patients (45.0% vs. 36.1%), which is consistent with literature as previous studies have reported that higher incidence of SCC was observed among Black males and females in comparison to white people^15^. Lung squamous cancer is more common in patients with a longer latency, regional stage and history of cervical squamous cell carcinoma. Thereby, some patients diagnosed with lung SCC may be metastatic from cervical cancer. In addition, patients with a history of chemo-radiation therapy also have a higher rate of SCC. A study^16^ shows that radiation penetrates epidermis sufficiently to cause irreversible DNA damage in cells located beneath the epidermis causing squamous cell carcinoma. Cervical cancer variables do not affect the stage of CC-NSCLC patients.

Chemotherapy and radiation were proven risk factors for the development of second malignancies among cancer survivors^17^. Traditionally, radiation exposure is considered as one of the most important risk factor for cancer development. Radiation and chemotherapy treatments have a crucial role in the treatment of early cervical cancer, decreasing the risk of cancer recurrence and improving survival, but such treatments are also associated to an increased risk of second malignancies after exposure, especially, in long-term smokers ^18–21^. However, no significant high standardized incidence ratios were observed among the radiation group in a large population-based study using SEER data. The authors explained that a half of the patients were none/unknown status of radiotherapy which could explained such results ^17^. In our population-based study using SEER data, significant high incidence ratios were observed in radiation group. With the improvement of radiotherapy technology, the application of Intensity-modulated radiation therapy (IMRT) and 3dimensional conformal radiation therapy (3D-CRT) has become more and more common ^22,23^. Our data cannot assess whether IMRT and 3D-CRT will increase the risk of second primary tumor. No significant SIR was detected in surgical treatment group.

Finally, our study has several limitations including notably the lack of biological data including PD-L1 status ^24–26^. We failed to collect this crucial data as biomarker analysis has not been universalized in clinical practice until very recent years. Consequently, further studies are warranted for targetable driver mutations and PD-L1 in CC-NSCL. Furthermore, our study did not include the interactions of all possible risk factors on cervical cancer patients. Furthermore, the main limitation of the SEER data, like any retrospective study of treatment effects, is the lack of randomness in treatment regimens, which leads to confounding factors, and result may be biased and interpreted with caution despite the use of PSM to remedy this defect.

